# Structural identification of individual helical amyloid filaments by integration of cryo-electron microscopy-derived maps in comparative morphometric atomic force microscopy image analysis

**DOI:** 10.1101/2021.10.19.464873

**Authors:** Liisa Lutter, Youssra K. Al-Hilaly, Christopher J. Serpell, Mick F. Tuite, Claude M. Wischik, Louise C. Serpell, Wei-Feng Xue

## Abstract

The presence of amyloid fibrils is a hallmark of more than 50 human disorders, including neurodegenerative diseases and systemic amyloidoses. A key unresolved challenge in understanding the involvement of amyloid in disease is to explain the relationship between individual structural polymorphs of amyloid fibrils, in potentially mixed populations, and the specific pathologies with which they are associated. Although cryo-electron microscopy (cryo-EM) and solid-state nuclear magnetic resonance (ssNMR) spectroscopy methods have been successfully employed in recent years to determine the structures of amyloid fibrils with high resolution detail, they rely on ensemble averaging of fibril structures in the entire sample or significant subpopulations. Here, we report a method for structural identification of individual fibril structures imaged by atomic force microscopy (AFM) by integration of high-resolution maps of amyloid fibrils determined by cryo-EM in comparative AFM image analysis. This approach was demonstrated using the hitherto structurally unresolved amyloid fibrils formed *in vitro* from a fragment of tau (297-391), termed ‘dGAE’. Our approach established unequivocally that dGAE amyloid fibrils bear no structural relationship to heparin-induced tau fibrils formed *in vitro*. Furthermore, our comparative analysis resulted in the prediction that dGAE fibrils are closely related structurally to the paired helical filaments (PHFs) isolated from Alzheimer’s disease (AD) brain tissue characterised by cryo-EM. These results show the utility of individual particle structural analysis using AFM, provide a workflow of how cryo-EM data can be incorporated into AFM image analysis and facilitate an integrated structural analysis of amyloid polymorphism.

## INTRODUCTION

Amyloid is a conformational state of polypeptides, which may be accessed by various natively folded or intrinsically disordered proteins (Iadanza et al., 2018). Conversion of protein and peptides into the amyloid state results in an aggregation-prone, β-sheet-rich fold (Eisenberg and Sawaya, 2017), followed by self-assembly into twisted fibrils via a nucleation-dependent mechanism, with numerous intermediate, self-propagating and coexisting species (Chuang et al., 2018). These assemblies, in some cases, are required for the normal physiological functioning of an organism, and in others lead to the formation of cytotoxic species that are a hallmark of neurodegenerative diseases and systemic build-up of misfolded proteins (Dobson, 2017). The connection between pathology and amyloid structures remains of particular interest. All amyloid fibrils share a cross-β core architecture comprised of extended stacks of β-strands that run perpendicular to the fibril axis, but which, importantly, can differ in cross-sectional folds and lateral organisation of protofilaments (Lutter et al., 2021). This diversity has been elucidated recently by the development of high-resolution structural analysis of fibrillar amyloid by cryo-electron microscopy (cryo-EM) and solid-state nuclear magnetic resonance (ssNMR) (Lutter et al., 2021; Willbold et al., 2021), as well as X-ray and electron diffraction of small amyloid peptide crystals (Sawaya et al., 2007, 2016). Proteins with an identical sequence are able to form different core or filament arrangements (Gremer et al., 2017; Wälti et al., 2016). Differences have also been demonstrated for *ex vivo* extracted and *in vitro* assembled fibrils (Bansal et al., 2021), with further differences arising from varying *in vitro* sample preparation conditions and techniques (Colvin et al., 2016; Schmidt et al., 2015). In particular, recent advances in cryo-EM have led to the elucidation of an array of structures of amyloid filaments formed by the tau protein (Shi et al., 2021). Tau is well known for its involvement in Alzheimer’s disease, but is also found in chronic traumatic encephalopathy, Pick’s disease and numerous other neurodegenerative diseases (Spillantini and Goedert, 2013). Cryo-EM studies have revealed that structures of tau fibrils may be disease-specific, although different diseases can also share the same tau folds (Hallinan et al., 2021; Shi et al., 2021). Importantly, cryo-EM-based structural models of heparin-induced tau filaments revealed significant differences between fibrils extracted from patient tissues and those generated synthetically *in vitro* (Arakhamia et al., 2020; Falcon et al., 2018b; Zhang et al., 2019)

Although cryo-EM and ssNMR methods can provide detailed structural information of the amyloid fibril core, both require measurements of samples with homogenous populations, or significantly populated subpopulations. ssNMR requires a ^15^N- and ^13^C-labeled homogenous sample, often achieved through rounds of consecutive seeding of fibrils with fresh monomer (Paravastu et al., 2009). However, seeding *ex vivo* extracted fibrils with recombinant monomer *in vitro* may not result in structures identical to those of the original *ex vivo* material (Koloteva-Levine et al., 2021; Lövestam et al., 2021). Non-seeded *in vitro* self-assembled fibril samples can also yield well-resolved spectra if the labelled starting material is mostly homogenous (Colvin et al., 2016), although *ex vivo* fibril structures cannot be studied without seeding with labelled monomer to produce fibrils amenable for ssNMR analysis. Cryo-EM can achieve high resolution structural models for significantly populated polymorphs of a heterogenous sample by classification of sub-populations. However, reconstruction requires averaging of tens to even hundreds of thousands of identical fibril segments. Therefore, it may not be possible to assess the polymorphic extent of the sample or to reconstruct polymorphs which are present in low numbers, but which may still be biologically relevant. Additionally, the analysis of intrafibrillar variation of structure and morphology of fibrils is limited due to the low signal-to-noise ratio from individual cryo-EM micrographs.

Atomic force microscopy (AFM) has recently emerged as a method capable of addressing amyloid structural polymorphism on an individual fibril level (e.g. Adamcik and Mezzenga, 2018; Aubrey et al., 2020). AFM is a technique in which the sample surface is probed with a sharp tip attached to a cantilever, creating 3-dimensional topographs at nanometre image resolution with a high signal-to-noise ratio. It enables the surfaces of individual amyloid fibrils to be reconstructed and the polymorphism landscape of the sample to be analysed on a single-molecule basis (Aubrey et al., 2020; Lutter et al., 2020). Therefore, AFM is highly complementary to cryo-EM and ssNMR, which rely on particle or ensemble averaging to produce reconstructions of fibril core structures. The application of AFM to structural biology has been hampered by the imaging artefact that stems from the physical finite size of the AFM probe, resulting in dilation and distortion of surface features. However, we have recently shown that a computational deconvolution approach is able to correct for this artefact, and the recovered surface sampling can result in an increased local image resolution. Furthermore, samples exhibiting helical symmetry, including amyloid fibrils, can be reconstructed as 3D models, facilitating structural analysis of the fibril surface envelopes (Lutter et al., 2020). AFM image deconvolution and 3D reconstruction of helical filaments can then be employed to carry out a quantitative mapping of amyloid fibril polymorphism at an individual fibril level (Aubrey et al., 2020). This deconvolution and reconstruction approach opens up the possibility of integrating high signal-to-noise individual filament 3D envelope structures from AFM data with high resolution ensemble-averaged core structural information available from cryo-EM and ssNMR methodologies. To date, AFM imaging of amyloid fibrils has been carried out alongside cryo-EM and ssNMR studies to determine fibril handedness or visualise the sample supra-structural morphology (Ghosh et al., 2021; Li et al., 2018; Röder et al., 2019). However, a fully integrated approach that links AFM data with cryo-EM derived structural maps has the potential to enhance the structural analysis of amyloid fibril polymorphism as AFM provides structural data on an individual fibril level, which is highly complementary to the averaged structural models from cryo-EM and ssNMR data. Nevertheless, there is currently no unifying workflow which would enable comparison of 3D-information from AFM surface envelope reconstructions with the structural maps provided by cryo-EM and ssNMR.

Here, we report a new approach to quantitative structural comparison of individual amyloid fibrils imaged by AFM with ensemble-averaged structural data derived from cryo-EM. This approach integrates AFM 3D envelope models of individual fibrils with cryo-EM maps and has been applied to the structural characterisation and identification of individual filaments formed by tau (297-391) known as ‘dGAE’ (Al-Hilaly et al., 2020), which has not yet been structurally resolved to high resolution by cryo-EM or by ssNMR. dGAE tau self-assembles spontaneously *in vitro* without the presence of anionic cofactors, such as heparin, to form filaments that share the height and cross-over characteristics observed in paired helical filaments (PHFs) isolated from Alzheimer’s disease tissue (Al-Hilaly et al., 2020). We compared the 3D surface envelope models of synthetic, *in vitro* assembled dGAE tau filaments constructed from previously published AFM image data with existing cryo-EM density maps of both *ex vivo* extracted tau fibrils and those formed *in vitro* by the addition of heparin, found in the Electron Microscopy Data Bank (EMDB). This analysis showed that the surface of dGAE amyloid fibrils closely resembles that of the cryo-EM density maps arising from some of the tauopathy tissue-extracted tau filaments, notably those of PHFs isolated from *ex vivo* Alzheimer’s disease tissue. Thus, our cryo-EM data-integrated AFM image analysis ruled out structural similarity of dGAE amyloid fibrils assembled *in vitro* to *in vitro* formed heparin-induced tau filaments, and has provided evidence that identified dGAE fibrils are structurally closely related to PHFs isolated from AD brain tissue. Individual fibril level structures in mixed polymorphic amyloid populations are likely to underpin properties such as overall fragmentation propensity, assembly kinetics and cytotoxic potential. Thus, by developing an integrative AFM image analysis, we reveal the potential for this form of AFM analysis to be used for identification of individual fibril structures in heterogeneous and polymorphic amyloid populations.

## RESULTS

### Three-dimensional reconstruction using AFM image data enabled structural analysis of individual dGAE tau amyloid filaments

AFM imaging of *in vitro* assembled dGAE filaments (Al-Hilaly et al., 2020) revealed a mixture of long individual fibrils and laterally associated, twisted bundles (**Fig 1a**). All fibrils and bundles showed a left-handed twist, with variation in the cross-over distance (*cod*, defined according to Supplementary **Fig S1a**). dGAE fibrils demonstrated a high propensity to bundle (**Fig 1a-b**). This may be indicative of surface properties of the dGAE protofilaments or fibrils, which favour the formation of large, twisted bundles of various widths. To analyse the structure of individual dGAE fibrils, 3D envelopes of individually selected single, non-bundled dGAE fibrils can be reconstructed from the AFM topographs using previously developed methods (Lutter et al., 2020). Here, to demonstrate this approach, a 3D surface envelope model of a single selected well-separated canonical dGAE fibril was reconstructed (**Fig 1c-d**). The central height profile of this dGAE filament showed peaks with the average height of 8.9 nm and grooves with the average height of 5.9 nm, with cross-over distances ranging from 43.1-58.6 nm, with an average of 47.7 nm (**Fig 1d**). These morphometric parameters are in line with the ranges expected of a typical non-bundled dGAE fibril observed by AFM (Al-Hilaly et al., 2020). During the 3D reconstruction workflow (Lutter et al., 2020), the fibril was determined to have 2-fold helical symmetry, suggesting that it is composed of 2-protofilaments arranged symmetrically about the central helical axis. The reconstructed 3D envelope model also enabled the evaluation of the cross-sectional area and shape, perpendicular to the fibril axis (defined in Supplementary **Fig S1a**), which was subsequently used in comparison with cryo-EM density maps of a range of different tau fibril structures.

**Figure 1.**
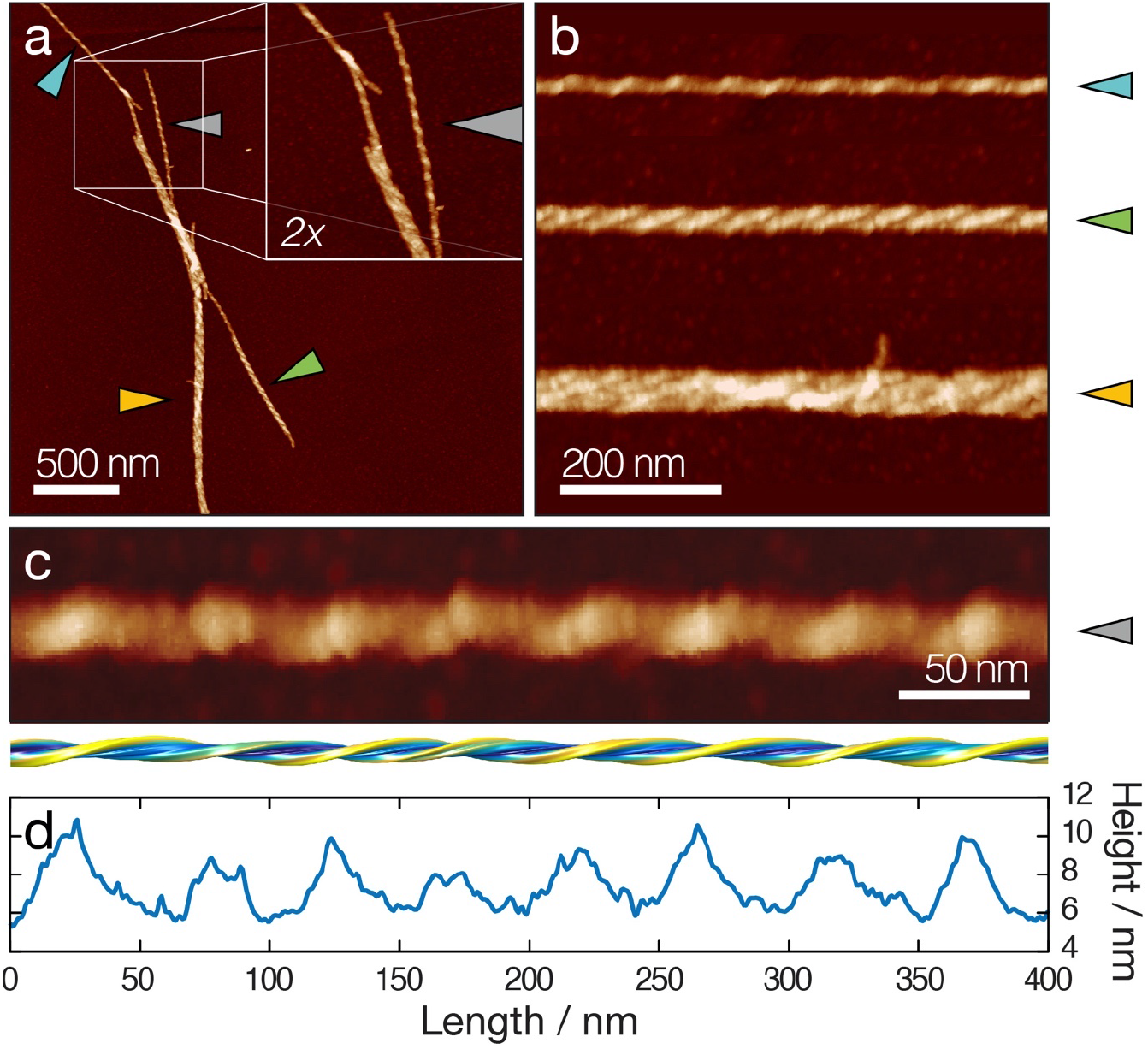
Analysis of individual dGAE filaments from a topographical AFM height image. (a) A typical AFM image of dGAE fibrils (Al-Hilaly et al., 2020) with an inset showing a 2x magnified view. Arrows point to the fibrils shown in (b) and (c). Scale bar represents 500 nm. (b) Digitally straightened images showing a typical dGAE fibril and typical fibril bundles. Scale bar represents 200 nm. (c) Digitally straightened AFM image of a canonical dGAE tau fibril together with a 3D reconstructed surface envelope model. Scale bar represents 50 nm. (d) The central line height profile of the fibril image shown in (c).

### Simulation of topographic images from volumetric cryo-EM maps allowed integration of cryo-EM data in comparative AFM image analysis

AFM imaging is able to generate image data in the form of 3D topographic raster scans, which are typically displayed as coloured 2D images where the pixel value represents surface height. As the images have a high signal-to-noise ratio, topographs of individual filaments can be traced and extracted from the surface scan data (**Fig 1**). In order to compare the 3D structural information from cryo-EM and AFM, simulated topographic AFM height images were generated from cryo-EM density maps (examples in **Fig 2**). This enabled the comparative analysis of the dGAE fibril imaged by AFM (**Fig 1c-d**) and cryo-EM data of patient derived tau fibrils and *in vitro* self-assembled heparin-induced tau fibrils.

**Figure 2.**
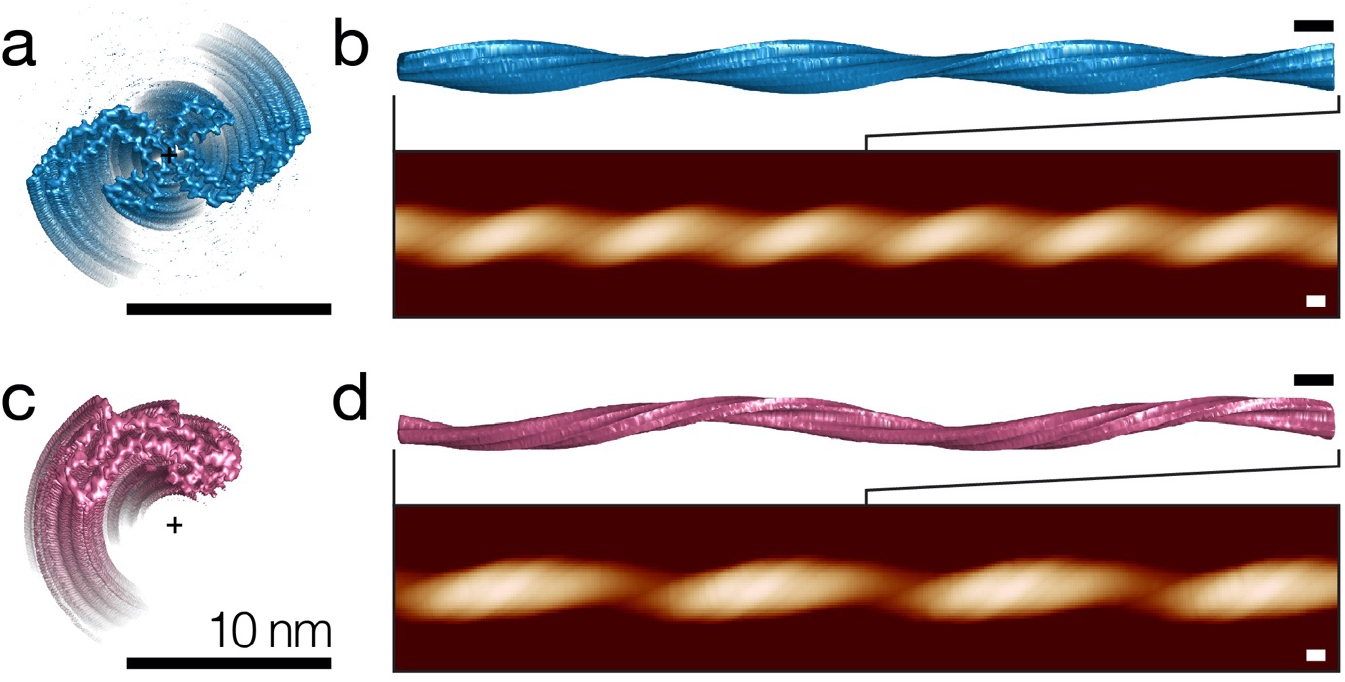
Simulation of topographic AFM height images from cryo-EM density maps. Cross-sectional views of axis-aligned cryo-EM density map iso-surfaces for (a) paired helical filaments (EMD-0259) and (c) ‘heparin-snake’ (EMD-4563) tau fibrils are shown with the position of the helical axis (+). Simulated topographic images of 500 nm fibril segments and the top view of extended 250 nm segments of iso-surfaces aligned to the simulated images are shown for PHF in (b) and ‘heparin-snake’ in (d). All scale bars represent 10 nm.

Volumetric cryo-EM maps of tau amyloid fibrils were first downloaded from the Electron Microscopy Data Bank, followed by extraction of the map iso-surface 3D coordinates at the iso-values specified by the authors of each cryo-EM entry. Where necessary, the individual maps were then denoised and the maps of the short fibril segments were extended by finding the helical axis of the map, followed by rotation and translation of multiple copies of the maps about their helical axes, using author reported twist and rise values, to a total map length equivalent to that of the AFM imaged dGAE fibril. Although helical twist and rise values are provided by the authors in experimental metadata, the position and angle of the fibril map screw-axis is unknown for all cryo-EM maps. Therefore, an iterative approach was used to find the rotation and translation parameters that provide the best alignment of the cryo-EM fibril maps to their respective helical axis positions (**Fig 2a, c**). In some cases, for example heparin-induced recombinant tau ‘snake’ fibrils, the resulting screw-axis position lies off the fibril centre-of-mass, resulting in a hollow centre (**Fig 2c)**, in line with previous analyses (Zhang et al., 2019). In order to simulate topographic AFM images of the extended iso-surfaces, a model of the AFM cantilever probe tip, which interacts with the filament surface, is then needed. The tip angle and geometry were estimated using the values provided by the manufacturer of the AFM probe used experimentally. The tip radius used in the simulations was matched to that used to image the dGAE fibril, which was estimated using AFM image data from the fibril (**Fig 1c-d**) according to a previous published algorithm (Lutter et al., 2020). A simulated AFM tip constructed from these geometry, angle, and radius parameters was then used to find the tip-sample contact points with the extended fibril iso-surface, to account for, and to reproduce, the tip-sample convolution effect in the simulated images. The sampling rate was also set to be equivalent to that of the experimental AFM data (**Fig 2b, d**). As both the tip radius and sampled pixel density of the simulated images match that of the individual fibril from the AFM image data, it was possible to make a direct comparison between the two.

### Simulated topographic AFM images from cryo-EM maps of helical fibrils enabled validation of the AFM based 3D reconstruction methodology

Prior to a full comparative analysis of the AFM derived 3D model of the dGAE fibril with simulated AFM images from published cryo-EM structural data of tau fibrils, the simulated AFM images were first used to validate the AFM-based 3D reconstruction methodology used to construct the dGAE fibril model (**Fig 1d**). In order to evaluate the 3D reconstruction method that we developed to reconstruct fibril 3D envelopes from AFM data and to determine their cross-sectional shapes and helical symmetries, reconstructions from simulated images of fibrils with known 3D structures were carried out, using cryo-EM maps of selected tau fibril structures (Supplementary **Fig S2** and **S3**). In this validation process, AFM topographs were simulated as described above, using the same pixel size and AFM tip radius as seen in the experimental AFM image of the dGAE tau fibril. Reconstruction of 3D envelopes was then carried out from these simulated images in the same way as from experimental AFM images, with the aim of comparing the final 3D reconstructions to the known fibril structures from which the input images were simulated. This allowed the validation of the reconstruction method regarding the accuracy of the cross-sectional shape, symmetry assignment, and positioning of the screw-axis, which the comparative image and morphometric analyses of dGAE tau fibrils requires. During the reconstruction workflow (Lutter et al., 2020), fibril symmetry is initially unknown so envelopes of fibrils with various symmetries, as well as an asymmetrically cross-sectioned fibril, were reconstructed from the simulated image data. The correct symmetry can then be identified by the angle of the twisting pattern on the fibril surface by comparison of topographs re-simulated from various symmetry models with the input topograph of the experimental or simulated image, in direct, as well as in Fourier space (Supplementary **Fig S2f** and **S3f**).

The cryo-EM density iso-surfaces for PHF (EMD-0259) and heparin-jagged (EMD-4565) fibrils were used to validate the 3D reconstruction method by comparison of the final 3D model with the known input structures. PHF and heparin-jagged maps were chosen to evaluate the reconstruction method, as the structures have C2 and C1 rotational symmetry, respectively, allowing the reconstruction of different symmetry assignments to be assessed. In addition, heparin-jagged has a helical axis that lies off the centre-of-mass of the core density. Evaluation of the approach demonstrated that the 3D reconstruction method can accurately and unambiguously identify both the C2 and C1 helical symmetries from fibril topographs (Supplementary **Fig S1f** and **S2f**). Furthermore, for PHF, the shape of the fibril cross-section closely matched that of the AFM tip-accessible cross-section from the original cryo-EM density map. The root-mean-square deviation (RMSD) between the tip-accessible PHF density map cross-section, which was obtained by tracing a circle with the radius equivalent to that of the AFM tip radius around the iso-surface cross-section to account for the topographic resolution imposed by the finite size of the AFM probe (Supplementary **Fig S1d**), and 3D reconstructed envelope (Supplementary **Fig S2e**) was 0.08 nm. For heparin-jagged fibril, with an off-centre screw-axis, cross-sectional RMSD was 0.23 nm (Supplementary **Fig S3e**), although the helical axis was correctly positioned to be lying off the fibril centre-of-mass (cross in Supplementary **Fig S3e**). These results validated the AFM based 3D reconstruction and simulation approach, although the RMSD values showed potential for improvement for the reconstruction workflow for asymmetric fibrils with off centre helical axes. More importantly, these results demonstrated that the approach correctly identified the symmetry and screw-axis position using known structural maps as input. This gave confidence that the 2-fold symmetry identified for the dGAE fibril from the experimental image data is accurate, indicating that it is composed of 2-protofilaments arranged symmetrically about a central helical axis, like the PHF from AD brain. Furthermore, as the RMSD between the known input and the 3D reconstruction was sub-Ångström for the 2-fold symmetric PHF example, the shape of dGAE fibril is also not likely to be significantly misrepresented by the 3D reconstruction process from the AFM data, although limitations can still potentially arise from the sample structure’s local mechanical stability and imaging conditions.

### Comparison of cryo-EM derived topographs with dGAE tau AFM data ruled out structural similarity with heparin-induced tau fibrils

Simulated topographic images from helical axis-aligned and lengthened cryo-EM density maps of *ex vivo* tissue-extracted and *in vitro* assembled heparin-induced tau filaments were subsequently prepared for comparison with the topographic AFM image of the canonical dGAE tau filament (**Fig 3**). This was achieved by carrying out topographic image simulations for all unique tau fibril cryo-EM maps available in the EMDB up to August 2021, which included structural maps of fibrils purified from tissues of patients with corticobasal degeneration (CBD), chronic traumatic encelopathy (CTE), Pick’s disease (PiD), Alzheimer’s disease (AD), as well as four fibril polymorphs formed *in vitro* by addition of heparin to recombinant tau monomers. In addition to the selected *ex vivo* fibril data, entries in the EMDB of near-identical structures to the selected PHFs and straight filaments (SFs) were also determined for filament structures from various other neurodegenerative diseases, including prion protein cerebral amyloid angiopathy (PrP-CAA) and primary age-related tauopathy (PART) (Hallinan et al., 2021; Shi et al., 2021). Where several near-identical structures have been reported for the same disease, the one with the highest map resolution was used. The helical twist of amyloid fibrils within a population of fibrils is variable even when the core structures are identical (Zhang et al., 2020), but the helical twist parameters presented in the population averaged cryo-EM maps represent only one of a continuum of possible values within a range, while AFM images report the full structural variation within an individual filament. To account for this difference in the information content, the periodicities of the simulated fibril images were optimised to maximise the normalised cross-correlation of the simulated and experimental fibril images by allowing the long fibril axis of the simulated image to contract or dilate within a range (see Methods). This allowed for slight adjustments of the simulated fibril periodicity, which aids the assessment of image similarity between experimental AFM data and simulated images derived from cryo-EM maps.

**Figure 3.**
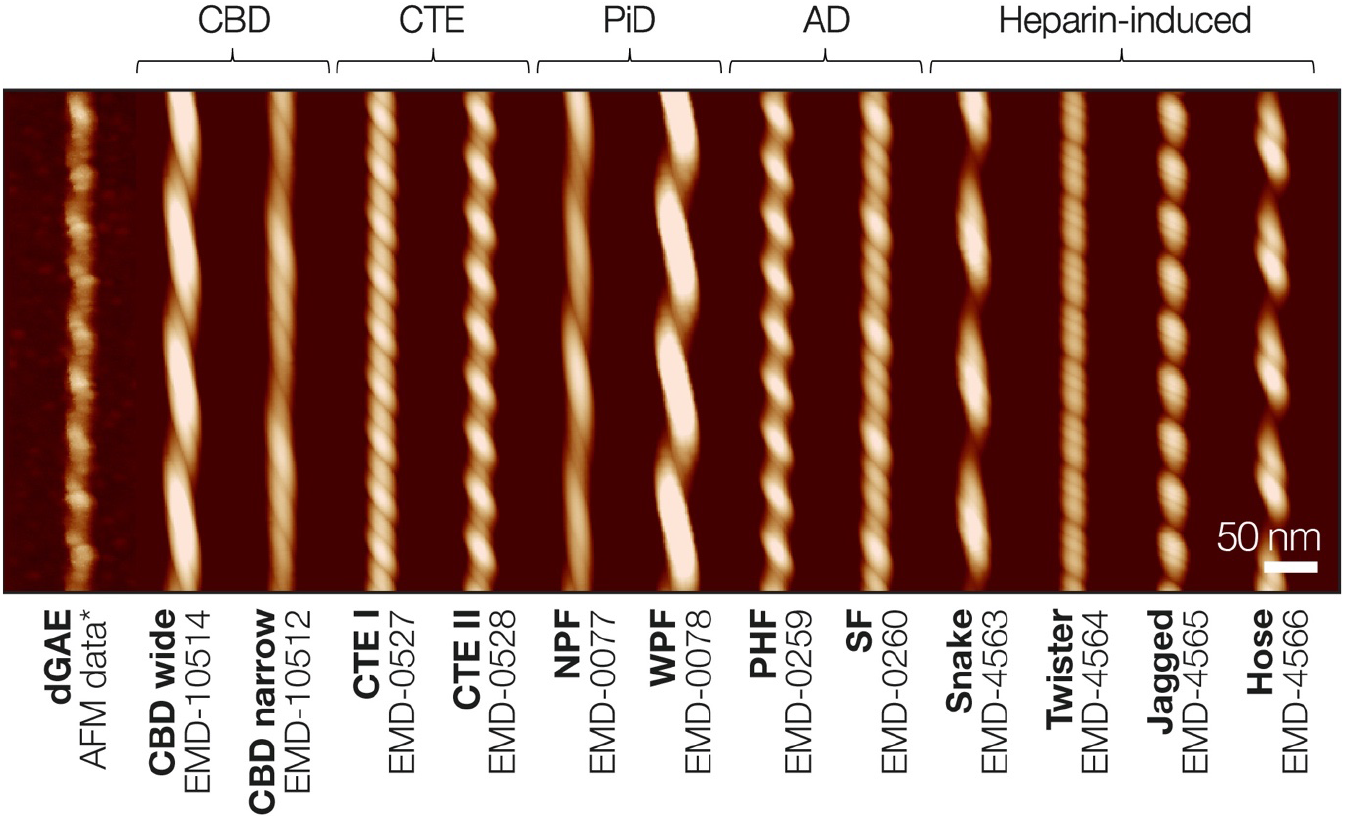
Comparison of the typical experimental AFM height image of dGAE tau fibril with topographical images of tau amyloid fibril polymorphs. The tau fibril polymorphs were simulated with identical tip (here estimated to be 11.7 nm) and sampling parameters using extended and axis-aligned cryo-EM map iso-surfaces. The ‘*’ indicate image data shown in **Fig 1c**. Length of the scale bar is 50 nm. The simulated images have been aligned to the same estimated helical axis height position as the experimental data image and, therefore, have comparable height colour coding. The EMDB accession codes of the cryo-EM density maps from which the topographs were simulated from are listed with the abbreviated names of the individual filaments. The sample origin is noted above the filaments’ images. The following abbreviations were used for disease diagnoses and filament labels: CBD – corticobasal degeneration, CTE – chronic traumatic encephalopathy, PiD – Pick’s disease, AD – Alzheimer’s disease, NPF – narrow Pick filament, WPF – wide Pick filament, PHF – paired helical filament, SF – straight filament.

The digitally straightened dGAE fibril (**Fig 1c-d**) is shown in **Fig 3** together with all of the simulated images of tau fibrils from published cryo-EM maps. All simulated tau fibrils are left-handed, as is the dGAE filament. As with any topographic AFM height image, the intensity of the colour also represents the height value in the simulated topographs. The comparison of fibril heights (colour intensity in **Fig 3)** immediately highlights the differences between the dGAE fibril and CBD wide filaments, as well as wide Pick filaments (WPF), both consisting of two intertwined protofilaments with larger cross-sectional areas compared to the dGAE fibril. However, CBD narrow filaments, narrow Pick filaments (NPF), and straight filaments (SF) from AD, which appear to be similar to dGAE in height, exhibit a difference in the repeating pattern compared to dGAE, due to their asymmetric cross-sections. These types of asymmetric cross-sections lead to a complex pattern of peaks and grooves along the length of the fibril, compared to the single visible peak and groove of the dGAE with 2-fold screw-symmetry, which in comparison is more similar to PHFs from AD. A similar pattern, although with a repeating unit consisting of a major and a minor peak, resulting from tip-sample contact with the rhombic cross-sections, are also seen on CTE type I and CTE type II filaments. Finally, the morphologies of all four heparin-induced tau fibrils differ from those of the other simulated images, and from the dGAE fibril image, due to their off-centre helical axis and asymmetric cross-sections. Although the dGAE fibril shown here matches the heparin-twister fibril well in height, the displayed repeating pattern differs substantially from the experimental dGAE fibril image due to differences in the cross-sectional shape and position of the screw-axis.

To quantify the similarity between the experimental dGAE fibril image and the simulated topographical AFM images derived from cryo-EM maps, a correlation distance based score (*d_img_*, see Methods) was calculated between the dGAE fibril image and each of the simulated images (Supplementary **Table S1**). Because the pixel sampling rate in the simulations was set to be identical to that of the experimental AFM image, each pixel value (height value) in the data image can be compared to the pixel value in a simulated image at the same coordinate location. The correlation distance score, *d_img_*, is then one minus the correlation coefficient between the data pixel values and the simulated pixel values, with smaller distance scores signifying more similar images. The *d_img_* score was calculated using the pixel value at the coordinates in the experimental AFM data image where the height values are above the estimated height of the fibril axis to minimise complex contributions from tip-sample convolution. The resulting *d_img_* scores (Supplementary **Table S1**) confirmed that the experimental dGAE fibril image showed closest similarity to the simulated AFM images derived from cryo-EM data of patient-tissue purified *ex vivo* PHF and CTE type II filaments.

### Quantitative and comparative morphometric analysis of dGAE tau amyloid fibrils with cryo-EM derived structural maps confirms their structural similarity to patient-derived tau structures

To investigate further whether dGAE tau amyloid fibrils are structurally similar to the tau amyloid purified from patient tissues, a quantitative and comparative morphometric analysis was performed next. First, the rotational cross-section of the dGAE tau fibril from the 3D model constructed using AFM data (**Fig 1c-d**), and those of the cryo-EM tau fibril density maps, were quantified in terms of their area and shape differences (cross-sectional area, *csa*, and cross-sectional difference, *csd*, respectively, defined according to Supplementary **Fig S1c-d**). For cryo-EM density maps, the rotational cross-sections were calculated by de-rotating the cross-sections along their helical axes, followed by tracing of the boundaries of the resulting 2D-maps with a circle of a radius equivalent to that of the AFM tip of the experimental dGAE fibril image. This procedure made the cross-sections from cryo-EM and AFM data comparable by accounting for the finite size of the tip during AFM imaging and thus estimating ‘tip-accessible’ cross-sections from the cryo-EM maps. A visual comparison of the dGAE fibril cross-section with CTE type II, PHF, and heparin-twister filament cross-sections (**Fig 4**) showed that although CTE II and PHF fibrils both display two-fold helical symmetry and show similarity in their simulated topographs to the dGAE fibril image data (**Fig 3**), the dimensions of the dGAE fibril model cross-section are smaller. However, although the area of the heparin-twister fibril cross-section better matched that of the dGAE fibril, it has an asymmetric cross-sectional shape with the helical axis positioned off the centre-of-mass, like other heparin-induced fibrils, which are dissimilar to the dGAE fibril with two-fold helical symmetry. Therefore, the differences in the helical symmetry and average cross-over distance were also taken into account in the comparative analysis, in addition to the cross-sectional difference and area parameters. The differences for all of these morphometric parameters between the dGAE tau fibril and patient tissue-extracted tau filament structures, or *in vitro* heparin-induced filaments, as determined by cryo-EM, were compiled and listed in Supplementary **Table S1**. These morphometric distance-based similarity scores, together with the image similarity score, *di_mg_*, allowed the calculation of a combined score of similarity, *d∑* (see Methods), between the dGAE fibril imaged by AFM and each of the unique previously determined cryo-EM tau fibril structural maps. This combined score enabled ranking of overall structural similarity (Supplementary **Table S1**), showing that the canonical dGAE fibril (**Fig 1c-d**) exhibits the highest overall structural similarity to PHF from Alzheimer’s disease, followed closely by CTE type II filaments, while unequivocally ruling out any close structural similarity to *in vitro* heparin-induced tau fibrils, as well as other disease-associated tau fibrils (**Fig 5**).

**Figure 4.**
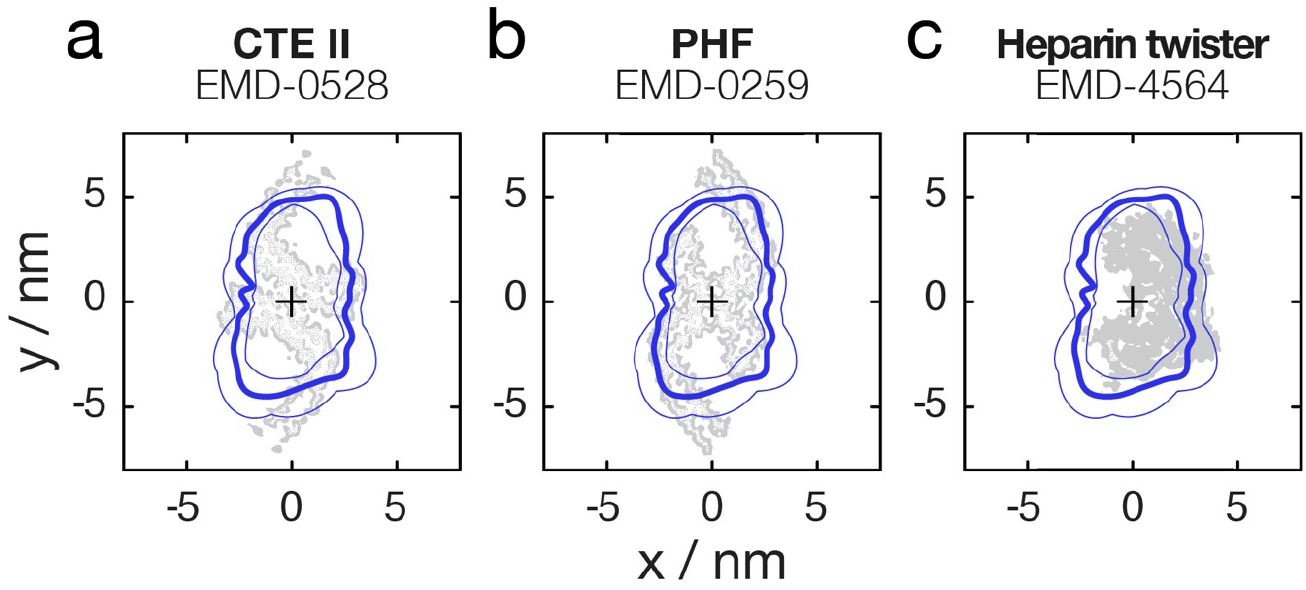
Comparison of the cross-section of the surface envelope model derived from the experimental AFM dGAE filament image and cryo-EM density maps cross-sections. The cross-sections for the CTE type II filaments (a) and PHF from AD patient tissue (b), as well as *in vitro* assembled heparin-induced ‘twister’ filaments (c) are shown. The average cross-section of the AFM image derived 3D model is shown as thick blue lines, and the structural variation observed along the fibril in the AFM image is indicated by the thin blue line denoting the 2.5 and 97.5 percentile bounds. The cryo-EM density cross-sections are shown in grey.

**Figure 5.**
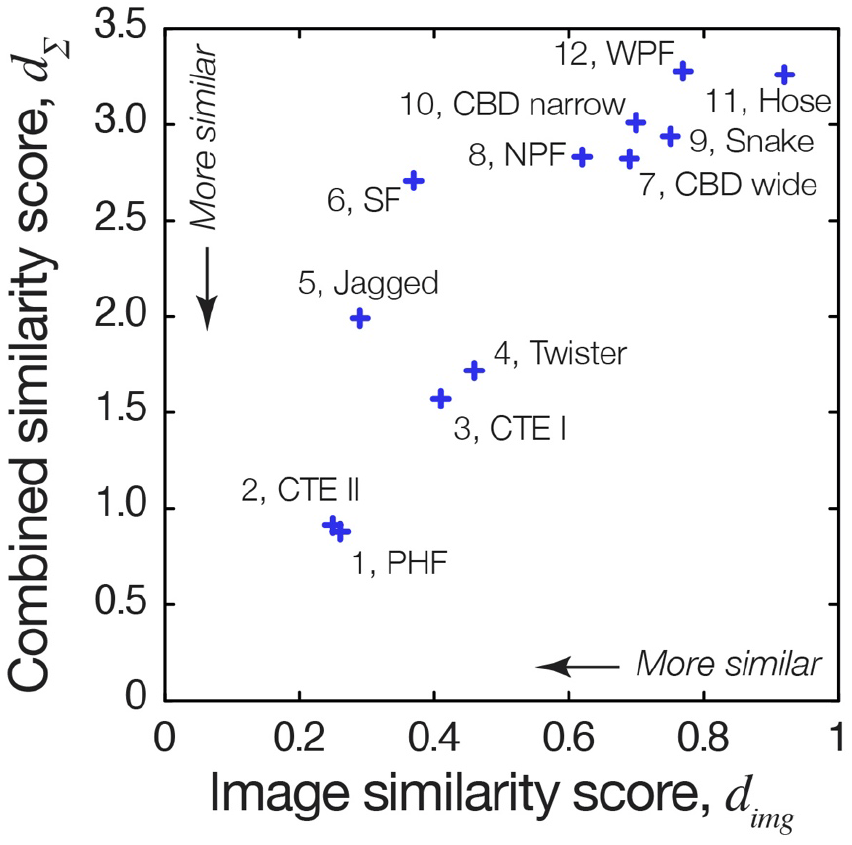
The image similarity score and the combined score of similarity between the dGAE fibril AFM image data and cryo-EM derived density maps of tau fibril polymorphs. Lower score denotes higher degree of similarity. The overall comparative rankings (from highest to lowest) based on the combined score of similarity (Supplementary **Table S1**) are labelled for each entry.

## DISCUSSION

Structural analysis of amyloid fibrils at the level of individual filaments is key to the understanding of the structural polymorphism displayed by amyloid assemblies, which in turn may relate to the varying physiological effects of amyloid *in vivo*. AFM imaging is complementary to high-resolution, ensemble-averaging techniques like cryo-EM, due to its capability for the structural analyses of single, individual molecules, e.g. membrane proteins (Heath et al., 2021), DNA (Pyne et al., 2021) and amyloid filaments (Aubrey et al., 2020), as well as their chemical signatures, response to force, interactions and dynamics (e.g. Maity and Lyubchenko, 2019; Ruggeri et al., 2015; Watanabe-Nakayama et al., 2016) Here using dGAE tau amyloid fibrils, we show that the 3D information from AFM and cryo-EM can be integrated with each other by simulation of topographic images from cryo-EM density maps or by 3D reconstruction of fibril surface envelopes from AFM topographic data. As exemplified by the analysis of the dGAE tau amyloid structure, these workflows enable direct comparison between experimental AFM images and the increasing numbers of cryo-EM data deposited in the EMDB. We demonstrate this approach by comparing high-resolution structural maps of various tau amyloid fibril cores determined by cryo-EM with a single AFM topographic scan of a sample of dGAE tau fibrils formed *in vitro* without the addition of anionic cofactors. In this case, the quantitative comparison of simulated topographs derived from the cryo-EM density maps with an experimental AFM fibril image of a canonical dGAE tau fibril shows that the structure of dGAE fibrils differs unambiguously from the structures of *in vitro* assembled heparin-induced recombinant tau fibrils. The analysis also predicts a close structural match between dGAE tau fibrils and both PHFs and CTE type II fibrils extracted from patient brain tissues.

The biological implications of our findings are significant because it is necessary to establish conditions of tau self-assembly *in vitro* that could result in the formation of disease-relevant fibril structures, in order to enable studies of these structures along a disease-relevant aggregation pathway, as well as to facilitate accessibility to physiologically relevant fibril structures for the development of aggregation modulators. Like dGAE fibrils, PHFs and CTE II fibrils also have 2-fold helical symmetry and have a similar cross-sectional shape. The cross-section of dGAE fibrils as estimated by the AFM derived 3D model is, however, smaller than those derived from CTE II fibrils and PHFs (**Fig 4**). This may be due to the physical interactions of the AFM probe tip with the fibril surface despite the low force applied, the deposition of the fibrils for AFM scanning in air or underlying molecular differences. However, the similarity between dGAE fibrils and PHFs as well as CTE type II filaments is also supported by sequence comparison. The dGAE tau sequence contains residues that correspond to the repeat regions 3 (R3) and 4 (R4) of the 4-repeat tau isoform, as well as 9 residues from the C-terminus of the repeat region 2 (R2), and thus it is expected the morphology of the fibrils would more closely resemble that of the fibrils which also contain R3 and R4 residues in their cores but few or none R1 or R2 residues. Such fibrils include PHFs and SFs AD filaments, as well as the two CTE filaments (Falcon et al., 2019, 2018b). In contrast, filaments from CBD, PiD, and heparin-induced recombinant tau also contain a larger numbers of residues from the R2 or R1 regions within their core structures (Falcon et al., 2018a; Zhang et al., 2020, 2019).

Although the sampling pixel density and tip-sample convolution effect are controlled for and matched to experimental values during the simulation of height images using cryo-EM maps, other factors arising from experimental imaging conditions such as the sample deposition process or residual monomeric or small aggregated species sticking onto the imaged fibril or the probe tip will affect the quality of experimental AFM data. Further improvements to the resolution and imaging parameters of the experimental dataset, either during data acquisition or by computational post-acquisition algorithms, could improve the accuracy of the quantitative comparison and the ranking of the similarity between experimental and simulated fibril images. The method presented here can also be extended to include ssNMR derived structural models by using atomic coordinates deposited to the Protein Data Bank instead of cryo-EM density maps for comparison. Thus, the approach presented here demonstrates the vast potential of integrated structural analysis that utilises the complementary 3D information acquired by different physical principles.

## METHODS

### AFM image data

The AFM image data analysed here was collected on a Bruker MultiMode 8 scanning probe microscope with a Nanoscope III controller, and is part of a previously published data set (Al-Hilaly et al., 2020). The images were acquired using the ScanAsyst in Air mode using Bruker Scan Asyst Air probes with nominal spring constant of 0.04 N/m. The AFM image shown in **Fig 1** represents topographical height scans 3 μm x 3 μm in size at a resolution of 2048 x 2048 pixels. The height image was flattened using Bruker Nanoscope Analysis software to remove tilt and bow. All further image analysis was carried out in Matlab.

### AFM image analysis

Fibrils were traced (Xue, 2014; Xue et al., 2009) and digitally straightened (Egelman, 1986) using an in-house application. The radius of the AFM tip during scanning was estimated by a least-squares fit of tip-sample contact-point model on the mean cross-section of a straightened fibril across its length, with tip radius as fitted parameter (Lutter et al., 2020). A tip model was then constructed from the tip radius, and the tip angle and geometry provided by the manufacturer. The same tip model was later used for simulating AFM topographs from cryo-EM data. The average fibril cross-over distance was found as the highest frequency peak from the Fourier transform of the fibril centre line height profile. The 3D reconstruction of the traced and straightened dGAE fibril was carried out as previously described (Lutter et al., 2020).

### Cryo-EM volumetric data pre-processing

Cryo-EM density maps were downloaded from the Electron Microscopy Data Bank (EMDB). The voxel-based volumetric Coulomb density map data were imported into Matlab where all subsequent processing and analysis was carried out. The helical rise, twist and handedness values were obtained from the information provided in the EMDB metadata of each entry. The iso-surface of a density map was found by applying the isovalue provided by the authors of each of the EMDB data entries. The iso-surface vertices were rotated and translated, where necessary, so that the long fibril axis runs along the z-axis. Where necessary, the density maps were denoised by finding the nearest neighbours for all iso-surface vertices and subsequently applying a threshold value to remove vertices with a significantly larger mean distance from neighbours. The number of nearest neighbours and threshold were changed according to the noise level of each map by manual inspection. All maps were then further denoised by finding the alpha-shape of the iso-surface using an alpha radius of twice the voxel size of the EM density map, while suppressing holes and retaining only the vertices that belong to the largest single region of the isosurface.

### Identifying and aligning the fibril screw-axis

The position of the fibril screw-axis was estimated by iterative optimisation of the rotation and translation parameters while maximising the overlap between a voxelised isosurface and the same isosurface rotated and translated about the Z-axis by applying the helical rise and twist values from the EMDB entry of each density map. Depending on the map features, rotation and translation parameters of the map isosurface coordinates relative to the screw-axis were optimised jointly or either rotation or translation was initially optimised separately for an initial estimate, followed by joint optimisation of all parameters.

### Simulation of AFM topographs from cryo-EM structural data

The axis-aligned iso-surface coordinates were extended to double the length of an experimental AFM image fibril to allow for periodic alignment. This was done by rotating and translating the iso-surface vertices along the long fibril axis (the z-axis in axis-aligned iso-surface) using the helical twist and rise values from the EMDB metadata, followed by a 90° rotation so that the filament axis aligned with the y-axis. The tip model with the radius specific to the experimentally imaged fibril was then used to find the contact points between the tip model and the fibril surface on an XY-grid constructed with the pixel size equivalent to that of the experimental AFM image. The periodicity pattern of the simulated fibril images was subsequently optimised to take into account possible structural variations in the twist of filaments by finding a possible best match to that of the experimental AFM fibril image within a plausible bounded range. This is achieved by an iterative optimisation of a parameter which dilates or contracts the simulated fibril image along the length of the fibril within a bounded interval of 0.5-2 times the nominal pixel size and maximises the cross-correlation between the transformed simulated image and the straightened experimental fibril image.

### Calculation of cross-sectional area and shape differences

Filament cross-sections perpendicular to the fibril axis were found for both the AFM 3D envelope model of the dGAE tau amyloid fibril and tau amyloid fibril cryo-EM density maps by untwisting the coordinates to the same 2D plane along the length of the fibril, using the twist and rise values provided in the EMDB metadata for cryo-EM models, or the analysis of periodicity for AFM imaged fibrils. For EM density maps, the tip accessible cross-section was estimated as the alpha shape of the cross-section, which was then found with the alpha radius equivalent to the tip radius of the fibril on the experimental AFM image. The alpha shape was then rotated by 1° intervals for a total of 360° and for each interval, RMSD to the cross-section of the experimental AFM fibril envelope was calculated. The rotation angle that resulted in the lowest RMSD indicated the best rotational alignment of the two cross-sections. The cross-sectional area or difference area were then calculated as defined in **Fig S1 c-d**.

### Similarity scores and comparative structural ranking

A score to quantify the similarity between the digitally straightened experimental AFM image of the dGAE fibril and simulated AFM images using cryo-EM maps of different polymorph of tau filaments found in the EMDB was defined as the correlation distance, *d_img_*, between the pixel height values, which is one minus the correlation coefficient, *r*, between the data pixel values and the simulated pixel values (**Eq. 1**)

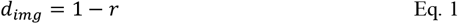

The *d_img_* score was calculated using only the pixel value at the coordinates in the experimental AFM data image where the height values are above the estimated height of the fibril axis to minimise complex contributions from tip-sample convolution. Scores to quantify the differences in the morphometric parameters (defined according to Supplementary **Fig S1**) helical symmetry, *sym*, twist handedness, *hnd*, average cross-over distance, *cod*, cross-sectional area, *csa* and cross-sectional difference area, *csd*, between each of the cryo-EM derived tau filament maps and the experimentally AFM-imaged dGAE fibril are defined as their individual Euclidian distances *d* (Aubrey et al., 2020), normalised to their observed maximum values, or the theoretical maximum value of 2 for *hnd*.

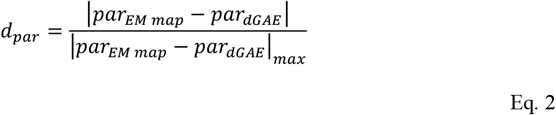

In **Eq. 2**, ‘*par*’ represents the parameters *sym, hnd, cod, csa* or *csd*. Defining of all the similarity scores using standardised distance measures means that lower numerical values indicate higher degree of similarity. The overall score of similarity, *d_Σ_*, was then defined as the sum of the individual image and morphometric similarity scores (**Eq. 3**).

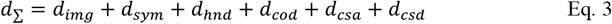

## ACKNOWLEDGMENTS

We thank Michel Goedert and Sjors Scheres for insightful discussions, and the members of the Xue group, the Kent Fungal Group and the Louise Serpell group for helpful comments throughout the preparation of this manuscript. This work was supported by funding from the Biotechnology and Biological Sciences Research Council (BBSRC), UK grant BB/S003312/1 (WFX and LCS), as well as Engineering and Physical Sciences Research Council (EPSRC), UK DTP grant EP/R513246/1 (LL). Research in the LCS lab, and YA-H is supported by funding from WisTa Laboratories Ltd (PAR1596).

## SUPPLEMENTARY FIGURES

**Figure S1.**
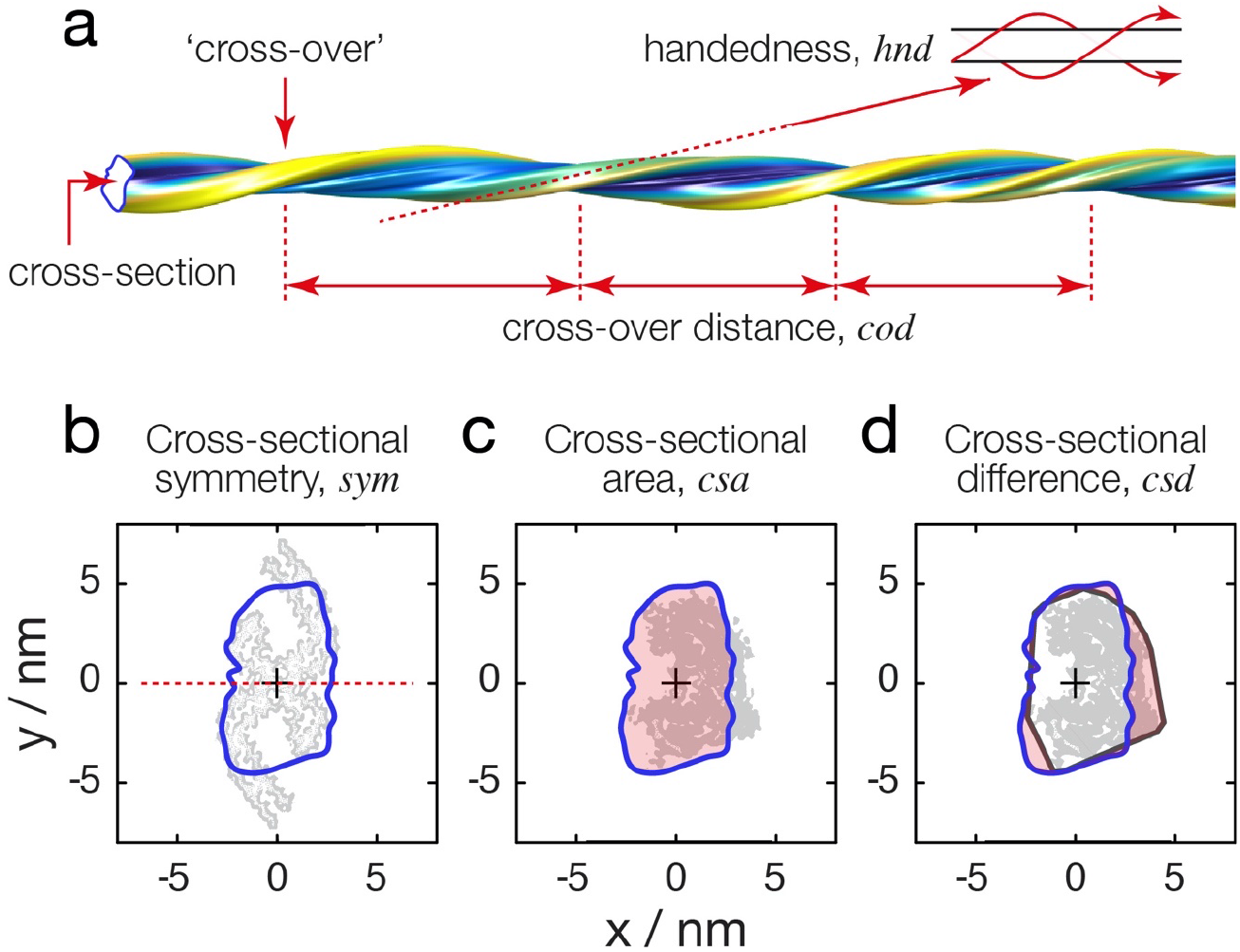
Schematic illustrations of the definitions of the morphometric parameters used for quantitative comparative structural analysis. (a) Schematic diagram illustrating the structural properties analysed, exemplified using a segment of the AFM derived dGAE fibril model shown in **Fig 1c**. The comparative morphometric parameters calculated by analysis of the cross-section are illustrated by the red line or areas in (b) for symmetry, (c) for cross-sectional area and (d) for cross-sectional difference area. Grey shapes show the cross-sections of cryo-EM density maps and the black line in (d) shows the tip-accessible cross-section, found by tracing a circle with the radius equivalent to that of the AFM tip radius around the cryo-EM density cross-section.

**Figure S2.**
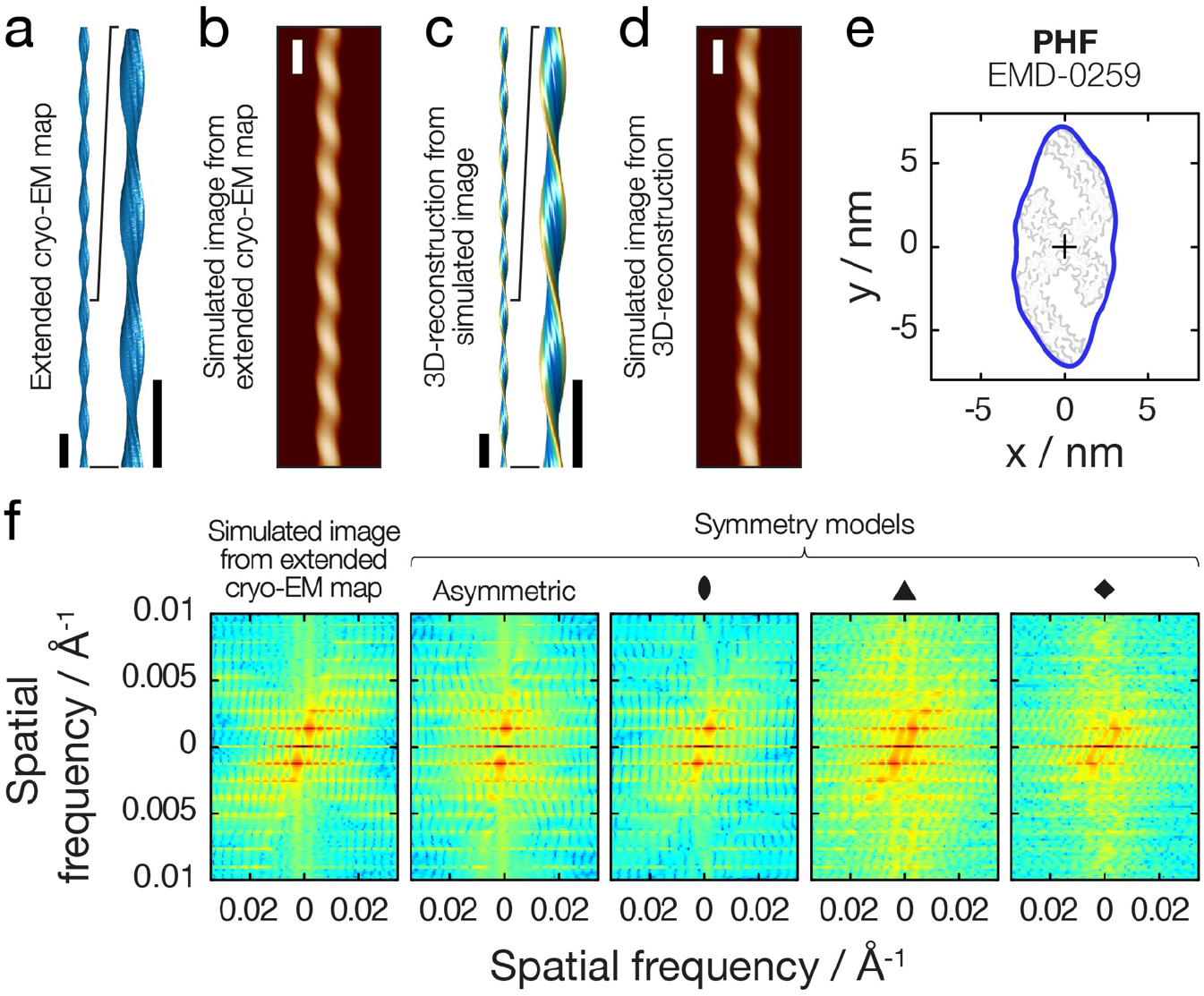
Validation of the 3D reconstruction and topograph simulation algorithms and workflow using a cryo-EM density map of PHF (EMD-0259). (a) Filament axis-aligned and lengthened cryo-EM density map of PHF. (b) Topographic AFM height image simulated from the density map in (a). (c) 3D surface envelope model reconstructed using the simulated image in (b). (d) Topographic AFM height image simulated from the 3D reconstructed model in (c). All scale bars represent 50 nm. (e) Comparison of the cross-sections of the EM density map (grey) and the AFM derived 3D model (blue). (f) Symmetry estimation during the 3D reconstruction process. As the symmetry is unknown, 3D reconstruction is performed with various symmetry estimates on the same AFM image data (here showing asymmetric, 2, 3, and 4-fold helical symmetries) and the 2D power spectra of the simulated topograph is compared to that of the original image (i.e. image shown in b). Here 2-fold symmetry gave the best match and was used for the final reconstruction.

**Figure S3.**
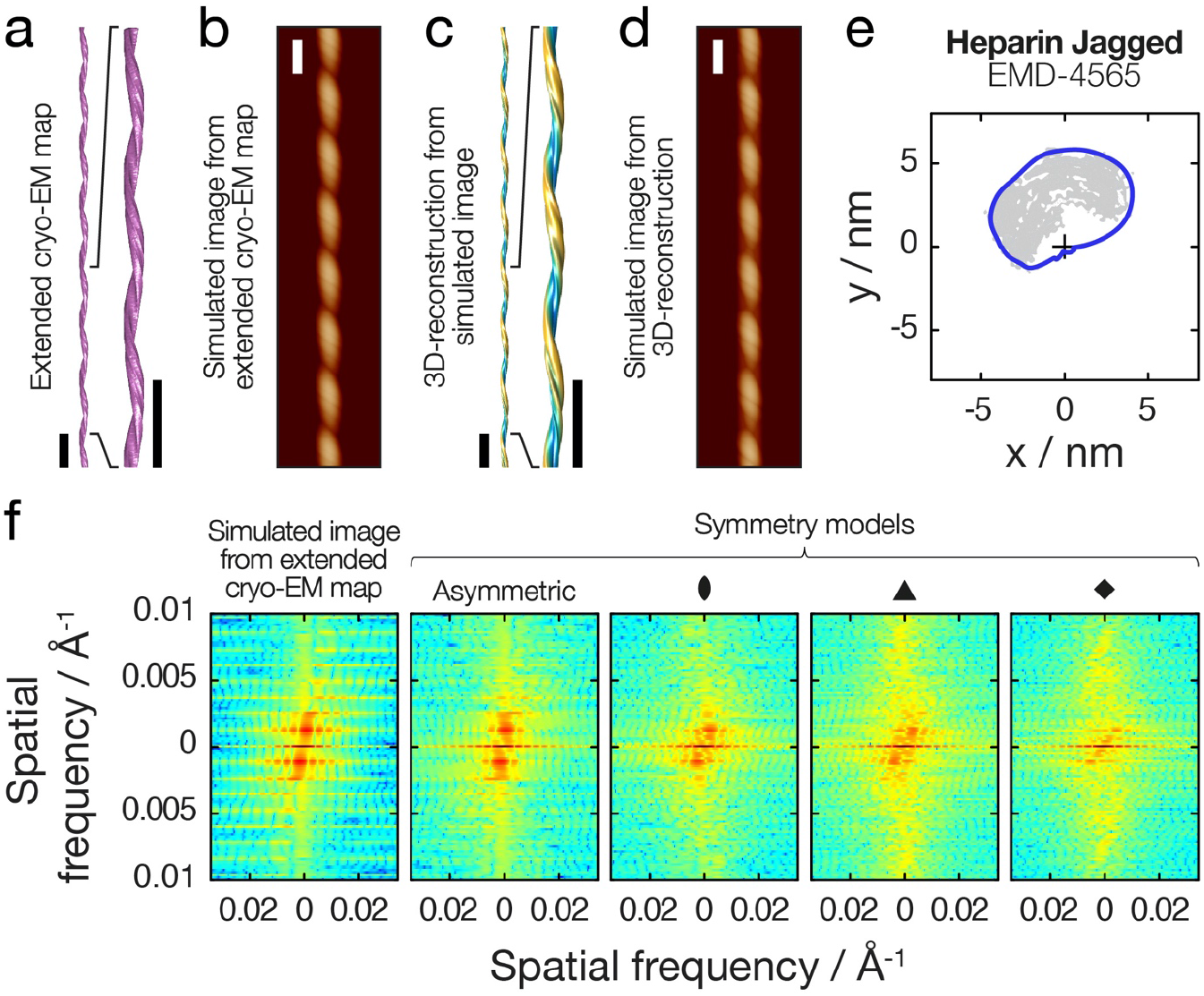
Validation of the 3D reconstruction and topograph simulation algorithms and workflow using a cryo-EM density map of a ‘heparin-jagged’ filament (EMD-4565). (a) Filament axis-aligned and lengthened cryo-EM density map of ‘heparin-jagged’. (b) Topographic AFM height image simulated from the density map in (a). (c) 3D surface envelope model reconstructed using the simulated image in (b). (d) Topographic AFM height image simulated from the 3D reconstructed model in (c). All scale bars represent 50 nm. (e) Comparison of the cross-sections of the EM density map (grey) and the AFM derived 3D model (blue). (f) Symmetry estimation during the 3D reconstruction process. As the symmetry is unknown, 3D reconstruction is performed with various symmetry estimates on the same AFM image data (here showing asymmetric, 2, 3, and 4-fold helical symmetries) and the 2D power spectra of the simulated topograph is compared to that of the original image (i.e. image shown in b). Here the asymmetric model gave the best match and was used for the final reconstruction

## SUPPLEMENTARY TABLE

**Table S1.** Quantitative comparison of image similarity and similarity in morphometric parameters between AFM image data of the dGAE fibril shown in **Fig 1c** and cryo-EM derived tau fibril maps using the distance measure-based scoring system described in the Methods section. The similarity rank is based on the combined score *d_Σ_*, from most similar with lowest *d_Σ_* value to least similar with highest *d_Σ_* value.

| *Filament type* | *sym* | *hnd*<sup>§</sup> | *cod* / nm | *csa* / nm<sup>2</sup> | *csd* / nm<sup>2</sup> | *d<sub>img</sub>* | *d<sub>sym</sub>* | *d<sub>hnd</sub>* | *d<sub>cod</sub>* | *d<sub>csa</sub>* | *d<sub>csd</sub>* | *d<sub>Σ</sub>* | Rank |
| --- | --- | --- | --- | --- | --- | --- | --- | --- | --- | --- | --- | --- | --- |
| dGAE, AFM data | 2 | -1 | 49.1<sup>†</sup> | 44.1 | 0.0 |  |  |  |  |  |  |  |  |
| CBD wide, EMD-10514 | 2 | -1 | 141.2 | 106.1 | 62.8 | 0.69 | 0 | 0 | 0.51 | 0.82 | 0.80 | 2.82 | 7 |
| CBD narrow, EMD-10512 | 1 | -1 | 203.9 | 54.8 | 24.3 | 0.70 | 1 | 0 | 0.86 | 0.14 | 0.31 | 3.01 | 10 |
| CTE type I, EMD-0527 | 2 | -1 | 71.1 | 83.5 | 40.2 | 0.41 | 0 | 0 | 0.12 | 0.52 | 0.51 | 1.57 | 3 |
| CTE type II, EMD-0528 | 2 | -1 | 71.1 | 64.4 | 21.3 | 0.25 | 0 | 0 | 0.12 | 0.27 | 0.27 | 0.91 | 2 |
| Narrow Pick's filament (NPF) , EMD-0077 | 1 | -1 | 229.4 | 39.6 | 11.5 | 0.62 | 1 | 0 | 1.00 | 0.06 | 0.15 | 2.83 | 8 |
| Wide Pick's filament (WPF) , EMD-0078 | 2 | -1 | 141.0 | 119.9 | 78.2 | 0.77 | 0 | 0 | 0.51 | 1.00 | 1.00 | 3.28 | 12 |
| Paired helical filament (PHF) , EMD-0259 | 2 | -1 | 77.6 | 61.3 | 18.4 | 0.26 | 0 | 0 | 0.16 | 0.23 | 0.23 | 0.88 | 1 |
| Straight filament (SF) , EMD-0260 | 1 | -1 | 164.8 | 70.5 | 27.9 | 0.37 | 1 | 0 | 0.64 | 0.35 | 0.36 | 2.71 | 6 |
| Heparin snake, EMD-4563 | 1 | -1 | 134.3 | 48.0 | 52.2 | 0.75 | 1 | 0 | 0.47 | 0.05 | 0.67 | 2.94 | 9 |
| Heparin twister, EMD-4564 | 1 | -1 | 50.1 | 37.3 | 12.9 | 0.46 | 1 | 0 | 0.01 | 0.09 | 0.16 | 1.72 | 4 |
| Heparin jagged, EMD-4565 | 1 | -1 | 83.3 | 45.2 | 38.8 | 0.29 | 1 | 0 | 0.19 | 0.01 | 0.50 | 1.99 | 5 |
| Heparin hose, EMD-4566 | 1 | -1 | 161.1 | 30.6 | 42.6 | 0.92 | 1 | 0 | 0.62 | 0.18 | 0.54 | 3.26 | 11 |

§. Left-hand twisted filaments are assigned a *hnd* value of -1 and right-hand twisted filaments are assigned a *hnd* value of 1
†. The mean cross-over distance of the selected dGAE fibril shown in **Fig 1c-d**. The overall comparative similarity rank is unchanged if the population mean *cod* value of dGAE fibrils of 72.7 nm (Al-Hilaly et al., 2020) is used instead.

